# Functional redundancy and crosstalk between flavodiiron proteins and NDH-1 in *Synechocystis* sp. PCC 6803

**DOI:** 10.1101/2019.12.23.886929

**Authors:** Lauri Nikkanen, Anita Santana Sánchez, Maria Ermakova, Matthias Rögner, Laurent Cournac, Yagut Allahverdiyeva

## Abstract

In oxygenic photosynthetic organisms excluding angiosperms, flavodiiron proteins (FDPs) catalyze light-dependent reduction of O_2_ to H_2_O. This alleviates electron pressure on the photosynthetic apparatus and protects it from photodamage. In *Synechocystis* sp. PCC 6803, four FDP isoforms function as hetero-oligomers of Flv1 and Flv3 and/or Flv2 and Flv4. An alternative electron transport pathway mediated by the NAD(P)H dehydrogenase-like complex (NDH-1) also contributes to redox hemostasis and the photoprotection of photosynthesis. Four NDH-1 types haven been characterized in cyanobacteria: NDH-1_1_ and NDH-1_2_, which function in respiration; and NDH-1_3_ and NDH-1_4_, which function in CO_2_ uptake. All four types are involved in cyclic electron transport. Along with single FDP mutants (Δ*flv1* and Δ*flv3*) and the double NDH-1 mutants (Δ*d1d2*, which is deficient in NDH-1_1,2_ and Δ*d3d4*, which is deficient in NDH-1_3,4_), we studied triple mutants lacking either one of Flv1 or Flv3, and NDH-1_1,2_ or NDH-1_3,4_. We show that the presence of either Flv1/3 or NDH-1_1,2_, but not NDH-1_3,4_, is indispensable for survival during changes in growth conditions from high CO_2_ /moderate light to low CO_2_ / high light. Our results suggest functional redundancy and crosstalk between FDPs and NDH-1_1,2_ under the studied conditions, and demonstrate that the functions of FDPs and NDH-1_1,2_ are dynamically coordinated for the efficient oxidation of PSI and for photoprotection under variable CO_2_ and light availability.

**One sentence summary:** Flavodiiron proteins and NDH-1 complex ensure survival of cyanobacterial cells by cooperatively safeguarding the photosynthetic apparatus against excessive reduction

## INTRODUCTION

Photosynthetic organisms have evolved a variety of different regulatory mechanisms which are important for the protection of the photosynthetic machinery during rapid changes in environmental conditions. Fluctuating light intensities present particular risk to the photosystems due to over-reduction of the photosynthetic electron transport chain (PETC), particularly Photosystem (PS) I (Allahverdiyeva et al., 2015; Tiwari et al., 2016; Shimakawa et al., 2016). To counter this, cyanobacteria, algae, and plants (excluding angiosperms) employ C-type flavodiiron proteins (FDPs) as a strong photoprotective electron sink, directing excess photosynthetic electrons from downstream of PSI to O_2_ (Helman et al. 2003, Allahverdiyeva et al., 2013; Gerotto et al., 2016; Ilik et al., 2017; Chaux et al., 2017; Jokel et al., 2018; Alboresi et al., 2019). This process is referred to as the Mehler-like reaction and, in contrast to the Mehler reaction (Mehler 1957), does not produce harmful reactive oxygen species (ROS) (Vicente et al., 2002; Brown et al., 2019).

The genome of the *β*-cyanobacterium *Synechocystis* sp. PCC 6803 (hereafter *Synechocystis*) encodes four isoforms of FDPs, namely Flv1-4, which function in O_2_ photoreduction mainly as hetero-oligomers consisting of either Flv1 and Flv3 or Flv2 and Flv4 (Zhang et al., 2004; Mustila et al., 2016; Santana-Sanchez et al., 2019). Recently, we showed that under air level [CO_2_], Flv1/3 and Flv2/4 hetero-oligomers function in a coordinated and interdependent manner: Flv1/3 stimulates strong, but transient O_2_ photoreduction upon the onset of illumination or during increases in light intensity, while Flv2/4 hetero-oligomers catalyze a slower and limited steady state reduction of O_2_, also using electrons downstream of PSI (Santana-Sanchez et al., 2019). In elevated [CO_2_] and in air [CO_2_] at alkaline pH 9, the *flv4-flv2* operon encoding Flv2, Flv4, and the Sll0218 proteins is downregulated (Zhang et al., 2009; Santana-Sanchez et al., 2019). Nevertheless, a low level of Flv1/3 in elevated [CO_2_] stimulates strong steady state O_2_ photoreduction, whereas in air [CO_2_] at pH 9, Flv1/3 is solely responsible for strong but transient O_2_ photoreduction (Santana-Sanchez et al., 2019). Whilst *in vitro* assays showed that homo-oligomers of recombinant Flv1, Flv3 and Flv4 can reduce O_2_ with NADH and/or NADPH (Vicente et al. 2002, Shimakawa et al. 2015, Brown et al. 2019), FNR and reduced Fd are yet to be considered as possible donors to FDPs. Moreover, the use of *Synechocystis* mutants solely overexpressing Flv1 or Flv3 clearly demonstrated that, in contrast to *in vitro* experiments, homo-oligomers of Flv3 or Flv1 are not involved in O_2_ photoreduction *in vivo* (Mustila et al. 2016). Thus, the electron donor to FDPs remains to be elucidated *in vivo*.

In cyanobacteria, photosynthetic and respiratory electron transfer reactions coexist on thylakoid membranes and are intricately linked, necessitating strict regulation (Mullineaux, 2014). The NAD(P)H dehydrogenase-like complex 1 (NDH-1, photosynthetic complex I) is mainly localized in thylakoid membranes and involved in several bioenergetic reactions including cyclic electron transfer around PSI (CET), respiration and CO_2_ acquisition via the carbon concentrating mechanism (CCM) (for reviews see Battchikova et al. 2011; Ma and Ogawa 2015; Burnap et al. 2015; Peltier et al. 2016). The NDH-1 complex is present as several forms with distinct physiological roles: the NDH-1_1_ type features the subunits NdhD1 and NdhF1 and functions as Complex I of the respiratory electron transfer chain (Zhang et al., 2004; He et al., 2015; Saura and Kaila, 2019); the NDH-1_2_ form features the NdhD2 subunit instead of NdhD1, but it remains unclear whether NDH-1_1_ and NDH-1_2_ serve different physiological functions (Peltier et al., 2016). The NDH-1_3_ and NDH-1_4_ forms that comprise low-Ci-inducible, high affinity CO_2_ acquisition subunits NdhD3, NdhF3, CupA, and CupS, or constitutive, low-affinity CO_2_ acquisition subunits NdhD4, NdhF4, and CupB, respectively, function in conversion of CO_2_ to HCO_3_^-^ (Ohkawa et al., 2000a,b; Shibata et al., 2001; Zhang et al., 2004; Burnap et al., 2013; Han et al., 2017; Schuller et al., 2019). This process is possibly enabled by alkaline pockets at the thylakoid membrane created by NDH-1 proton pumping (Kaplan and Reinhold, 1999). All four NDH-1 types catalyze CET from ferredoxin (Fd) to plastoquinone (PQ), which is coupled to the pumping of 4H^+^/2e^-^ into the thylakoid lumen (He et al., 2015; Laughlin et al., 2019; Schuller et al., 2019; Saura and Kaila, 2019). However, for NDH-1_3,4_ this is to a lesser extent than for NDH-1_1,2_ (Bernat et al., 2011).

In the current study, we aim to elucidate how the electron transport pathways mediated by FDPs and NDH-1 cooperate to allow the maintenance of redox poise between the PETC, respiration, CCM, and CO_2_ fixation in the Calvin-Benson-Bassham cycle (CBB) under variable light and C_i_ availability. To this end, we employed both biophysical and biochemical methods to characterize various *Synechocystis* mutant strains with combined deficiencies of both FDP and NDH-1 pathways. Our results provide convincing evidence for the presence of either Flv1/3 or NDH-1_1,2_, but not NDH-1_3,4_, being indispensable for the survival of *Synechocystis* cells under transitions from high [CO_2_] conditions to the combined stress conditions of air [CO_2_] and high light. We show that a dynamically coordinated and cooperative function of Flv1/3 and NDH-1_1,2_ is required for the photoprotection of the photosynthetic apparatus of *Synechocystis* cells and discuss the molecular mechanisms involved.

## RESULTS

### Simultaneous inactivation of Flv1/3 and NDH-1_1,2_ is lethal upon shift from high to air [CO_2_] and high light

In order to examine whether the simultaneous inactivation of NDH-1 and FDPs has an adverse effect on cell survival, we monitored the growth of *Synechocystis* wild-type (WT) and various mutant strains lacking (i) either Flv1 or Flv3 (Δ*flv1* and Δ*flv3*, respectively, Helman et al., 2003), (ii) both NdhD1 and NdhD2 (Δ*d1d2,* deficient in NDH-1_1_ and NDH-1_2_, Ohkawa et al., 2000), (iii) both NdhD3 and NdhD4 (Δ*d3d4,* deficient in NDH-1_3_ and NDH-1_4_, Ohkawa et al., 2000), as well as triple mutants with combined deficiencies of both pathways (Δ*flv1 d1d2*, Δ*flv3 d1d2*, Δ*flv1 d3d4*, Δ*flv3 d3d4*) in conditions of differing CO_2_ availability and light intensity. The triple mutants exhibited similar or slightly slower growth compared to the WT and other mutant strains under high [CO_2_] (3% CO_2_) and moderate light (ML, intensity of 50 μmol photons m^-2^s^-1^ PAR) (Fig. 1A). We also detected no substantial difference in growth when the 3% CO_2_/ML-pre-grown cells were diluted (OD_750_=0.1) and shifted to air [CO_2_] conditions under constant light intensity (Fig. 1B, Fig. S1A, Fig. S1C). In contrast, when cultures pre-grown in 3% [CO_2_]/ML were diluted (OD_750_=0.1) and subjected simultaneously to air [CO_2_] and high light stress conditions (HL, intensity of 220 μmol photons m^-2^s^-1^), the growth of both Δ*flv3 d1d2* and Δ*flv1 d1d2* strains was completely inhibited (Fig. 1D). Importantly, Δ*flv1*, Δ*flv3* and Δ*d1d2* strains demonstrated growth similar to WT cells under air [CO_2_]/HL conditions.

**Figure 1.**
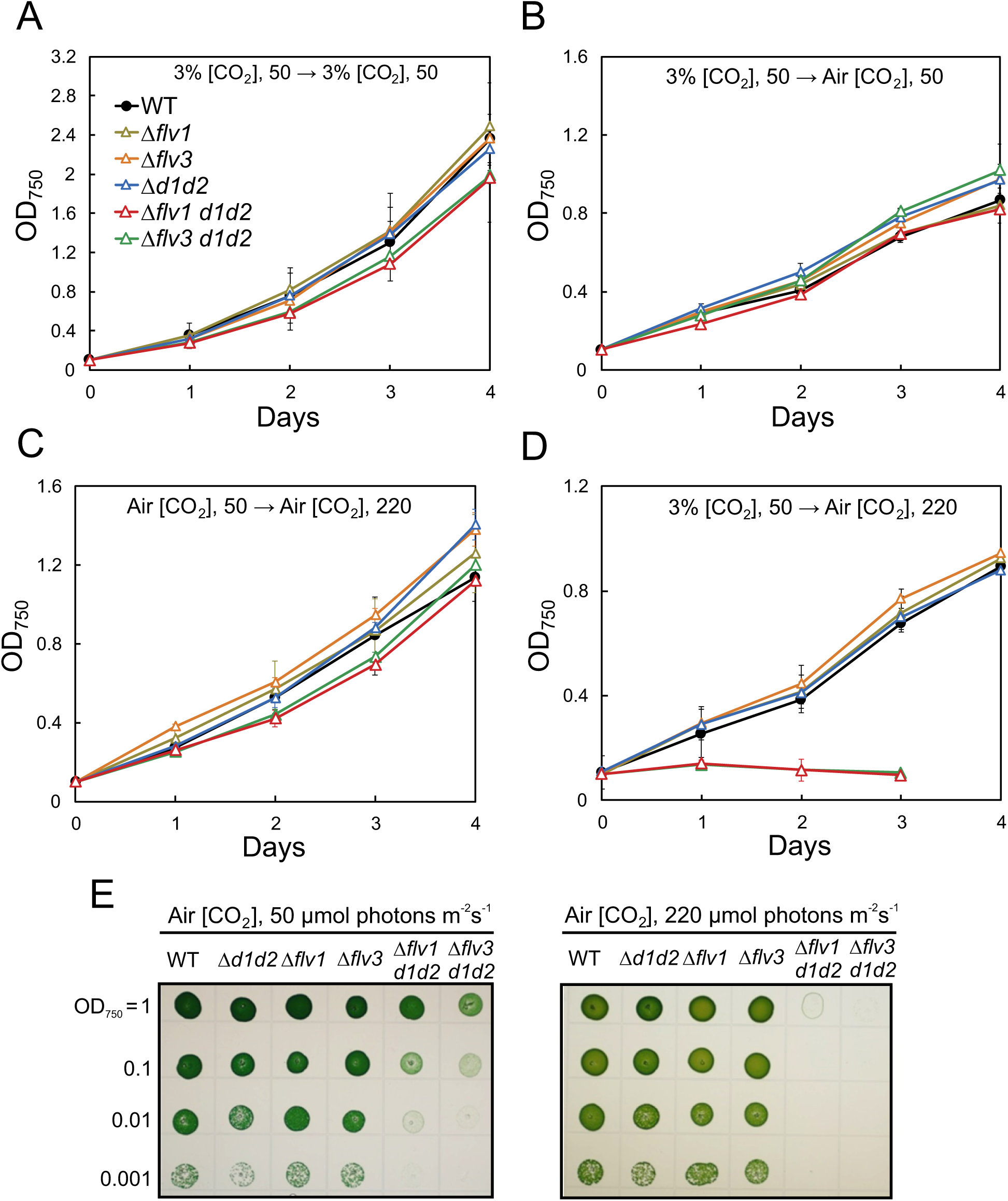
Growth of WT and mutant strains under different experimental conditions. All experimental cultures were started from pre-cultures adjusted to OD_750_=0.1. (A) Cells were pre-cultivated under 3% [CO_2_] / ML (50 μmol photons m^-2^s^-1^), adjusted to OD_750_=0.1 and grown in the same conditions. (**B**) Cells were pre-cultivated under 3% [CO_2_]/ML and shifted to air [CO_2_]/ML. (C) Cells were pre-cultivated under air [CO_2_]/ML and shifted to air [CO_2_]/HL 220 μmol photons m^-2^s^-1^). (**D**) Cells were pre-cultivated under 3% [CO_2_]/ML and shifted to air [CO_2_]/HL. The values presented in A–D are means of three independent biological replicates ±SE. (**E**) Growth of WT and mutant cells on agar plates containing BG-11. Cells grown under 3% [CO_2_] / ML were harvested and spotted on BG-11 agar plates in ten-fold dilution series starting at OD_750_=1, incubation was under air [CO_2_] /ML and air [CO_2_] /HL for 7 days. See also Supplemental Fig. S1.

We next examined whether pre-adaptation to air [CO_2_] would rescue the growth arrest. When cells previously adapted to air [CO_2_] /ML were diluted (OD_750_=0.1) and shifted to air [CO_2_] /HL, no differences in growth patterns between the strains were detected (Fig. 1C). Importantly, the triple mutants lacking either Flv1 or Flv3 and NDH-1_3,4_ (Δ*flv1 d3d4* and Δ*flv3 d3d4* strains) did not show a lethal phenotype after a shift from 3% [CO_2_]/ML to air [CO_2_]/HL, although a somewhat reduced growth was observed compared to the control strains (Fig. S1B, Fig. S1D). This suggests that, under the studied conditions, crosstalk between Flv1/3 and NDH-1_3,4_ complexes is not as crucial as with NDH-1_1,2_. Our attempts to create M55/Δ*flv1* and M55/Δ*flv3* double mutants lacking either Flv1 or Flv3 and the central membrane component of the NDH-1 complex, NdhB, were both unsuccessful. We were unable to obtain any M55/Δ*flv1* colonies and the M55/Δ*flv3* mutant strain demonstrated poor growth and was not segregated, suggesting that the absence of both Flv1/3 and NDH-1, disrupts essential cell metabolism.

We also studied the growth of the mutant strains on agar plates containing BG-11 (Fig. 1E). For these experiments, 3% CO_2_/ML-grown cells were diluted and grown on plates either under air [CO_2_] / ML or air [CO_2_] / HL conditions for 7 days. The Δ*flv1* and Δ*flv3* and Δ*d1d2* mutants did not exhibit any visible differences in growth capacity under these conditions in comparison to the WT. In contrast, the growth of the Δ*flv1 d1d2* and Δ*flv3 d1d2* triple mutants was strongly reduced under air [CO_2_] / ML, while no growth was detected under air [CO_2_] / HL (Fig. 1E). The Δ*flv1 d3d4* and Δ*flv3 d3d4* triple mutants demonstrated a slow growth phenotype, similarly to that previously reported for Δ*d3d4* on agar plates (Ohkawa et al. 2000b).

As the growth phenotypes of Δ*flv1* and Δ*flv3,* Δ*flv1 d1d2* and Δ*flv3 d1d2* were similar under all conditions, we only included Δ*flv3* and Δ*flv3 d1d2* in the majority of subsequent experiments.

### Either Flv1/3 or NDH-1_1,2_ is required for the oxidation of PSI during sudden increases in light intensity

The lethal phenotype observed upon the shift to air [CO_2_] / HL which was caused by the simultaneous impairment of the Mehler-like reaction catalyzed by Flv1/3 (Helman et al. 2003, Allahverdiyeva et al. 2011, 2013) and by NDH-1_1,2_ complex mediated respiration and CET (Ohkawa et al., 2000b, Bernat et al. 2011) suggests a functional redundancy between these electron transport pathways. To further investigate this possibility, we used membrane inlet mass spectrometry (MIMS) to examine the kinetics of O_2_ and CO_2_ exchange upon HL-illumination of dark-adapted WT and mutant cells grown for four days in air [CO_2_] / ML. In order to distinguish O_2_ uptake from photosynthetic gross O_2_ evolution under illumination, we enriched cell suspensions with ^18^O_2_ prior to measurements. In dark-adapted WT cells, a transient peak in O_2_ uptake occurs during the first minute of illumination (Fig. 2A). We showed recently that this transient peak is attributed to the Mehler-like reaction catalyzed predominantly by Flv1/3 hetero-oligomers, while Flv2/4 hetero-oligomers mainly contribute to steady-state light-induced O_2_ reduction in an interdependent manner (Santana-Sanchez et al., 2019). In Δ*flv3* cells, light-induced O_2_ reduction was almost abolished and only a slight impairment of the rate of photosynthetic gross O_2_ evolution was observed in comparison to WT cells (Fig. 2B). This result is in agreement with previous studies (Helman et al., 2003; Allahverdiyeva et al., 2013; Santana-Sanchez et al., 2019).

**Figure 2.**
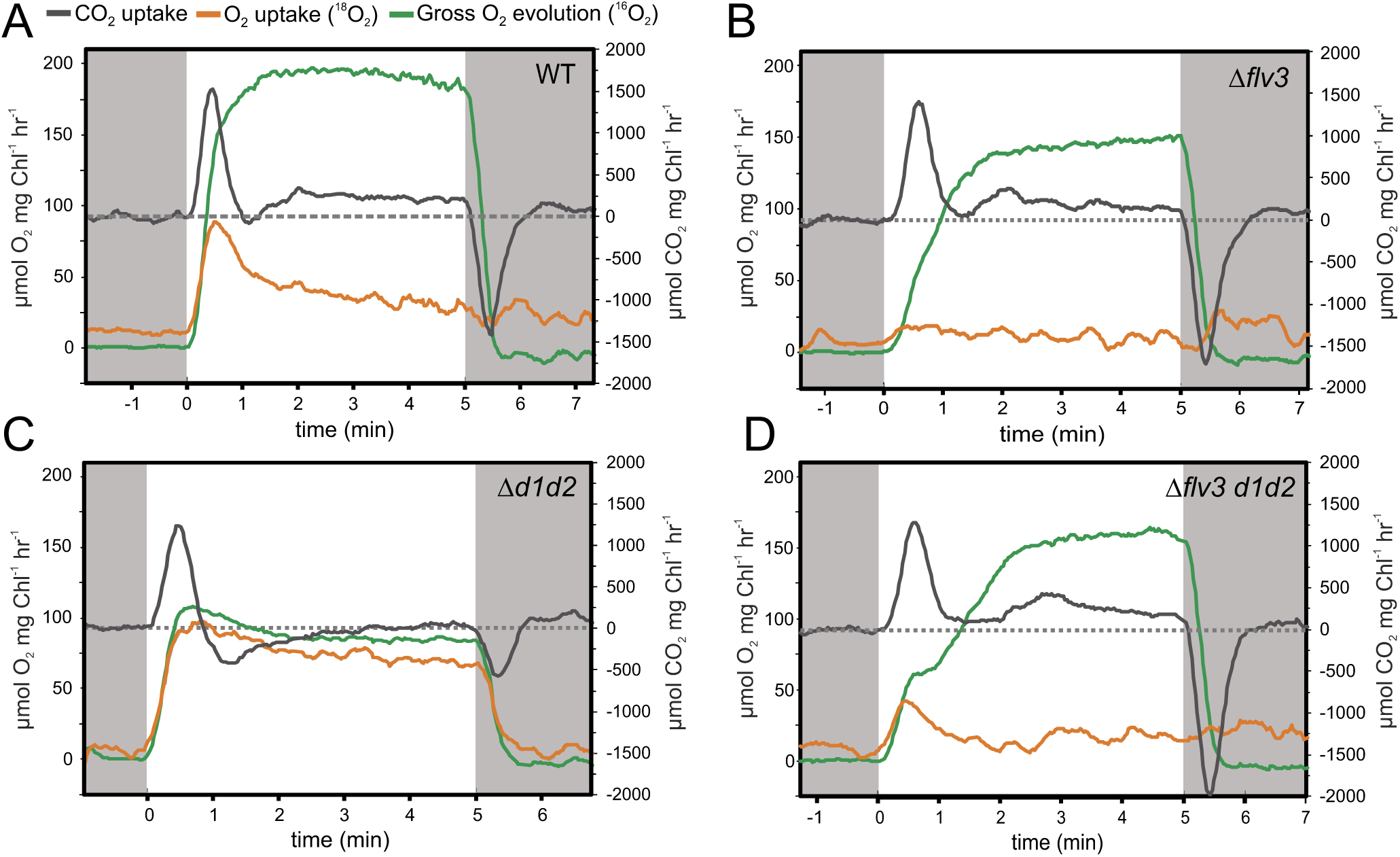
O_2_ and CO_2_ exchange rates in WT and mutant strains. Cells were grown in air [CO_2_] / ML for 4 days, after which the cells were harvested and Chl *a* concentration adjusted to 10 μg ml^-1^ with fresh BG-11. Cells were dark-adapted for 15 min, and gas exchange was monitored by membrane inlet mass spectrometry (MIMS) over a 5 min illumination period with 500 μmol photons m^-2^s^-1^ of white actinic light. Prior to the measurements, samples were supplemented with ^18^O_2_ at an equivalent concentration to ^16^O_2_ to distinguish O_2_ uptake from O_2_ evolution, and with 1.5 mM NaHCO_3_. The dashed line in each panel indicates the compensation point of CO_2_ fixation (uptake rate equals respiratory rate). The experiment was repeated with three independent biological replicates, of which representative measurements are shown.

In contrast to the WT, Δ*d1d2* cells lacked a fast decay phase of light-induced O_2_ uptake during the first minute of illumination, demonstrating sustained O_2_ photoreduction at high levels (c.a. 75–100 µmol O_2_ mg Chl^-1^ hr^-1^) throughout the illumination period. Meanwhile, gross O_2_ evolution was diminished nearly two-fold compared to the WT (Fig. 2C). Thus, sustained O_2_ photoreduction in Δ*d1d2* is not due to increased electron flow from PSII, but most likely due to increased activity of FDPs. Indeed, Δ*flv3 d1d2* mutant cells showed only minor light-induced O_2_ uptake transiently during the first minute of illumination (Fig. 2D). The Δ*flv3 d1d2* cells demonstrated slower induction of photosynthetic O_2_ evolution following two-component kinetics, and slightly impaired steady-state gross O_2_ evolution compared to WT (Fig. 2D).

The initial peak in CO_2_ uptake rate at the onset of illumination likely reflects activation of the CCM (Liran et al., 2018). Accordingly, this initial peak is absent in the M55 mutant deficient in NDH-1_3,4_ in addition to NDH-1_1,2_, as well as in WT cells grown at pH 6 (Fig. S2). No salient impairment of CCM was observed in any of the studied strains under air [CO_2_] / ML, with CO_2_ uptake rates peaking at c.a. 1.5 mM CO_2_ mg Chl^-1^ hr^-1^. In WT as well as in Δ*flv3*, CO_2_ fixation is then induced and the respiratory compensation point surpassed after ∼1 min of illumination (Fig. 2A-B). In Δ*d1d2*, however, induction of CO_2_ fixation is severely delayed, and the cells only reached the compensation point at the end of the 5 min illumination period (Fig. 2C). Intriguingly, although a slight delay was observed also in Δ*flv3 d1d2* induction of CO_2_ fixation was largely recovered in the triple mutant (Fig. 2D).

These MIMS results indicated that combined deficiency of Flv1/3 and NDH-1_1,2_ under standard growth conditions, air [CO_2_] / ML, slightly perturbs photosynthetic electron transfer and the redox poise between the PETC and cytosolic sink reactions. Next, we examined the ability of the mutant strains to adjust their photosynthetic activity to variable light conditions by measuring Chl *a* fluorescence simultaneously with P700 redox changes in conditions where light intensity periodically fluctuated between low (25 μmol photons m^-2^s^-1^, LL) and high irradiance (530 μmol photons m^-2^s^-1^, HL). Cells lacking Flv3 suffered from transient PSI acceptor side limitation Y(NA) upon sudden increases in light intensity, resulting in delayed oxidation of PSI (Fig. 3A), which is in line with previous studies (Helman et al. 2003, Allahverdiyeva et al. 2013). During subsequent 1 min cycles of LL/HL, however, the ability of Δ*flv3* to oxidize PSI in HL improved, suggesting that cells were able to acclimate to the fluctuating light condition via a compensatory mechanism that is distinct from Flv1/3 hetero-oligomers, such as *e*.*g*. NDH-1 mediated electron transport.

**Figure 3.**
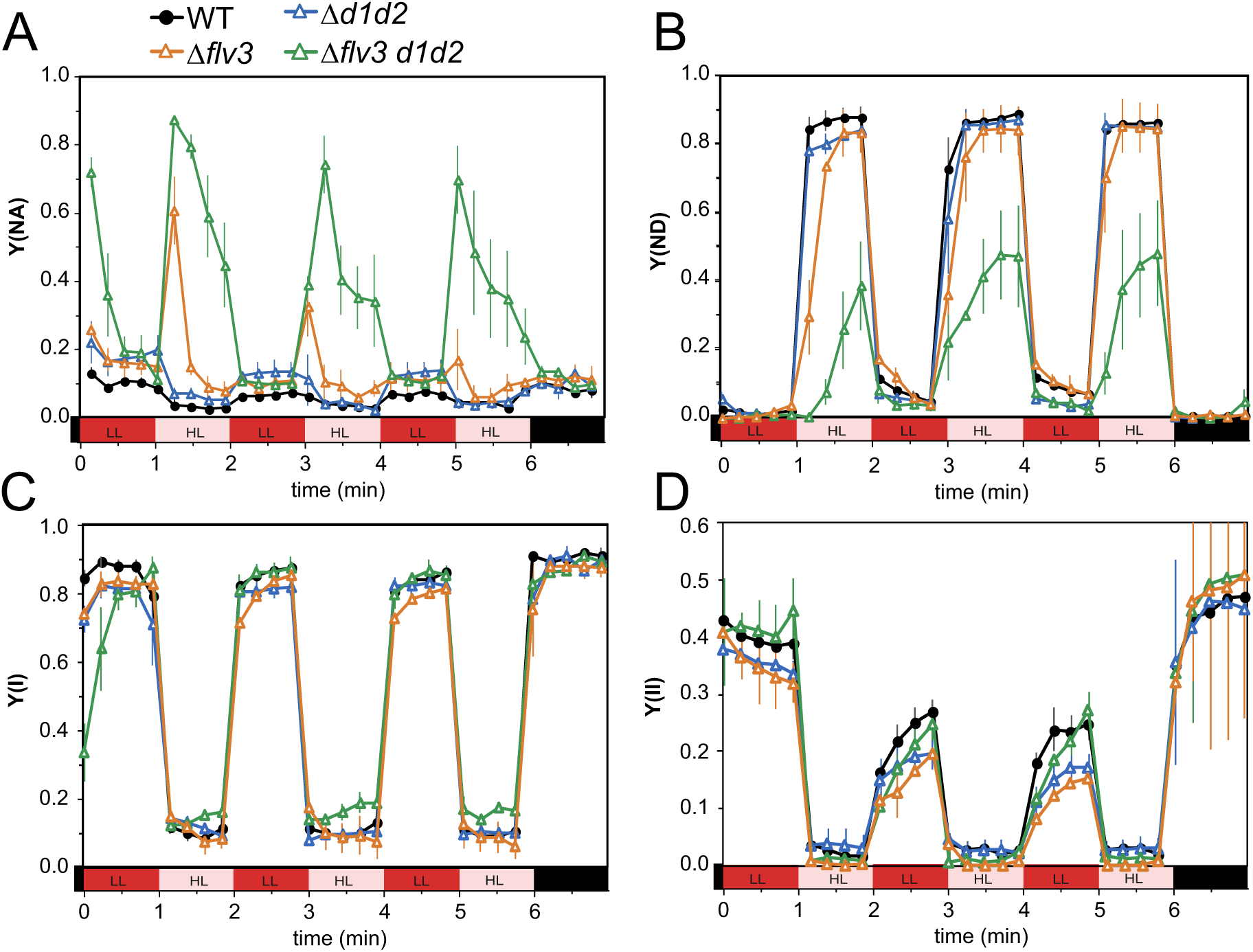
Photosynthetic parameters in WT and mutants strains in fluctuating light intensities. (**A**) Acceptor side limitation of PSI, Y(NA), (**B**) donor side limitation of PSI, Y(ND), as well as (**C**) effective yield of PSI, Y(I) were calculated from absorbance difference changes at 875-830 nm according to (Klughammer and Schreiber, 2008). Chl *a* fluorescence changes were measured simultaneously with PSI redox changes and used to calculate (**D**) effective yield of PSII. Cells were grown in air [CO_2_] / ML for 4 days, after which the cells were harvested and Chl *a* concentration adjusted to 10 μg ml^-1^ with fresh BG-11. Cells were dark-adapted for 10 min, and then illuminated with red actinic light intensity alternating between 25 and 530 μmol photons m^-2^s^-1^ in 1 min periods. Saturating pulses (500 ms of 5000 μmol photons m^-2^s^-1^) were provided every 15 seconds. The values shown are averages from 3–4 independent biological replicates ±SE.

We detected no acceptor-side limitation during the HL phases in Δ*d1d2* (Fig. 3A) as PSI was oxidized similarly to the WT (Fig. 3B). Interestingly, however, Δ*d1d2* cells exhibited slightly elevated acceptor-side limitation during the LL phases of the experiment (Fig. 3A), suggesting a role for NDH-1_1,2_ in maintaining photosynthetic redox poise in light-limited conditions. In the triple mutant strain Δ*flv3 d1d2*, transitions from low to high irradiance as well as from dark to light caused severe limitation on the acceptor side of PSI, resulting in an inability to oxidize PSI during periodic 1 min HL illumination. Unlike the Δ*flv3* mutant, Δ*flv3 d1d2* did not show improvement in the PSI acceptor side limitation during subsequent cycles. This implies a compensating activity of NDH-1 in Δ*flv3* mutant under the studied conditions. Interestingly, diminished donor side limitation in Δ*flv3 d1d2* resulted in slightly elevated effective yield of PSI during the HL phases (Fig. 3C). This could indicate the attenuation of pH-dependent limitation of electron transfer at Cyt *b*_6_*f*, due to impairment of proton motive force (*pmf*) generation in CET and the Mehler-like reaction. The effective yield of PSII was slightly decreased in HL in Δ*flv3 and* Δ*d1d2,* but not in Δ*flv3 d1d2* (Fig.

The differences in photosynthetic electron transport reported above may also be contingent on altered redox states of the NADP^+^/ NADPH pool. Therefore, in order to investigate the effect of Flv3 and/or NDH-1_1,2_ deficiency on NADP^+^/NADPH redox kinetics, we recorded NADPH fluorescence changes from dark-adapted cells simultaneously with Chl *a* fluorescence in conditions where actinic light intensity fluctuated between 25 (LL) and 530 μmol photons m^-2^s^-1^ (HL) similarly to Fig. 3. Upon dark-to-LL transitions, NADPH rapidly accumulated close to a maximal amount in all strains. Re-oxidation of NADPH then occurred in WT and Δ*flv3d1d2* cells, while in Δ*flv3* a slower oxidation phase preceded a transient re-reduction phase during the first minute of illumination (Fig. 4A-B). Strong reduction of the NADP^+^ pool was also detected in Δ*flv3* during the second HL phase of the experiment. In Δ*d1d2* cells, very little oxidation of the NADPH pool occurred during illumination, even at HL-LL transitions (Fig. 4C). Chl *a* fluorescence also remained at an elevated level (Fig. 4C), suggesting a reduced PQ pool. This was possibly due to delayed activation of CO_2_ fixation in the CBB cycle (Fig. 2C), and impaired CET or dark respiration. In contrast, NADPH was strongly oxidized in Δ*flv3 d1d2* cells at HL-LL transitions (Fig. 4D), but was predominantly reduced in dark-adapted cells (Fig. 4D). Upon cessation of illumination, oxidation of the NADPH pool was followed by transient re-reduction after c.a. 10 s. As observed previously (Schreiber and Klughammer, 2009; Holland et al., 2015), the re-reduction peak coincided with a secondary post-illumination rise in Chl *a* fluorescence (PIFR), while oxidation of NADPH paralleled an initial PIFR. The post-illumination re-reduction of NADP+ was diminished and both Chl *a* PIFR peaks were missing in Δ*d1d2* (Fig. 4C). Therefore, the PIFR peaks likely originate from cyclic and respiratory electron transport via NDH-1_1,2_ (Holland et al., 2015). Both Chl PIFR peaks were however observed in Δ*flv3 d1d2* cells, indicating involvement of a complementary pathway, possibly via succinate dehydrogenase (Cooley and Vermaas, 2001), or a CET pathway dependent on PGR5 and PGRL1-LIKE (Yeremenko et al., 2005; Dann and Leister, 2019). Finally, we monitored NADPH redox kinetics under conditions mimicking the MIMS experiments, where strong O_2_ photoreduction occurs in WT and Δ*d1d2* during the first minute of dark-to-high light transition (Fig. 2). No substantial differences were observed between the strains, as close to maximal reduction of NADP^+^ was obtained within ∼0.25 s (Fig. 4F) and maintained throughout a 40 s illumination period (Fig. 4E). This argues against NADPH being a primary electron donor to either Flv1/3 or NDH-1_1,2_.

**Figure 4.**
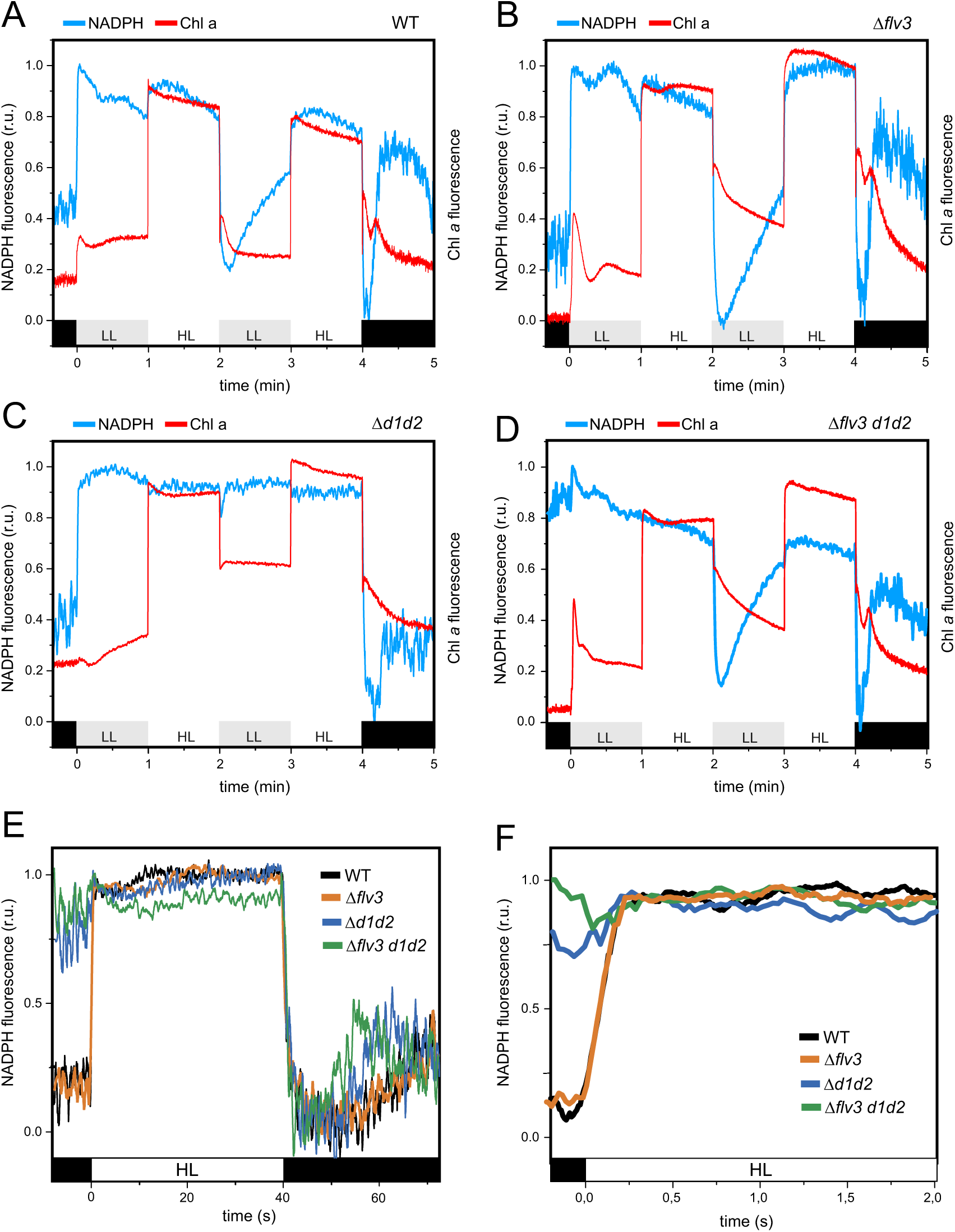
NADPH and Chl *a* fluorescence under fluctuating light in WT and mutant strains. WT (**A**), Δ*flv3* (**B**), Δ*d1d2* (**C**), and Δ*flv3 d1d2* cells (**D**) were grown in air [CO_2_] / ML for 4 days, after which the cells were harvested and Chl *a* concentration adjusted to 5 μg ml^-1^ with fresh BG-11. Cells were dark-adapted for 10 min, and then subjected to illumination with red actinic light alternating between 25 and 530 μmol photons m^-2^s^-1^ in 1 min periods. NADPH redox changes were monitored with the Dual-PAM 100 spectrophotometer and its 9-AA/NADPH accessory module by measuring fluorescence changes between 420 and 580 nm induced by excitation at 365 nm, as described by (Schreiber and Klughammer, 2009) and (Kauny and Setif, 2014). Chl *a* fluorescence was recorded simultaneously. (E) NADPH redox changes during illumination of dark-adapted cells with 530 μmol photons m^-2^s^-1^. Other experimental details are as in A-D. (**F**) shows a magnification of the first 2 s of illumination in (**E**). The values are normalized according to minimum and maximum fluorescence changes at the onset and cessation of illumination, respectively. The experiments were repeated with three independent biological replicates, of which representative measurements are shown. r.u. = relative units.

### Redox status of PSI donor and acceptor side electron carriers and buildup of *pmf* during dark-to-light transitions depend on both Flv1/3 and NDH-1_1,2_

To elucidate the molecular mechanism behind the photosynthetic phenotypes of the studied mutant strains, we next utilized a DUAL-Klas-NIR 100 spectrophotometer to distinguish between redox changes of plastocyanin (PC), P700, and ferredoxin (Fd) (Klughammer and Schreiber, 2016; Schreiber, 2017; Setif et al., 2019) upon exposure of dark-adapted cells to high irradiance (503 μmol photons m^-2^s^-1^). Upon illumination of WT cells, rapid oxidation of P700 and PC occurred, followed by transient reduction of PC, P700, as well as Fd after ∼0.2 s (Fig. 5A). Re-oxidation of all three electron carriers then ensued after ∼0.5 s. In Δ*flv3* cells P700 and PC remained mostly reduced after the initial oxidation transient until ∼3 s, when another transient and partial re-oxidation peak of P700 and PC occurred at ∼8 s (Fig. 5B). This was followed by re-reduction of P700, similarly to that observed recently by Bulychev et al. (2018). After 15 s in light, P700 and PC became gradually oxidized. In contrast to WT, Fd remained reduced for several seconds in light, and was only slowly re-oxidized over the 30 s illumination period (Fig. 5B). These observations indicate the importance of Flv1/3 as an electron sink to O_2_, accepting electrons presumably from reduced Fd (after ∼0.5 s in light). In Δ*d1d2*, redox changes of PC, P700 and Fd were similar to WT, except that the reduction of P700^+^ following initial oxidation as well as reduction of Fd occurred already after ∼50 ms (Fig. 5C). In Δ*flv3 d1d2*, re-reduction of P700^+^ and reduction of Fd also occurred already after ∼50 ms (Fig. 5D), suggesting involvement of NDH-1_1,2_ as an acceptor of electrons from Fd at that stage. Fd was reduced more quickly in Δ*flv3 d1d2* than in any of the other strains and was slowly re-oxidized over 30 s (Fig. 5D), likely due to shortage of electron acceptors (Fig. 3A). In summary, these NIR-spectroscopic measurements revealed that Flv1/3 and NDH-1_1,2_ control the redox poise between PSI, PC, and primary electron acceptor Fd during specific time frames at transitions to high irradiance. The results are congruent with Fd functioning as the electron donor to both NDH-1_1,2_ and Flv1/3.

**Figure 5.**
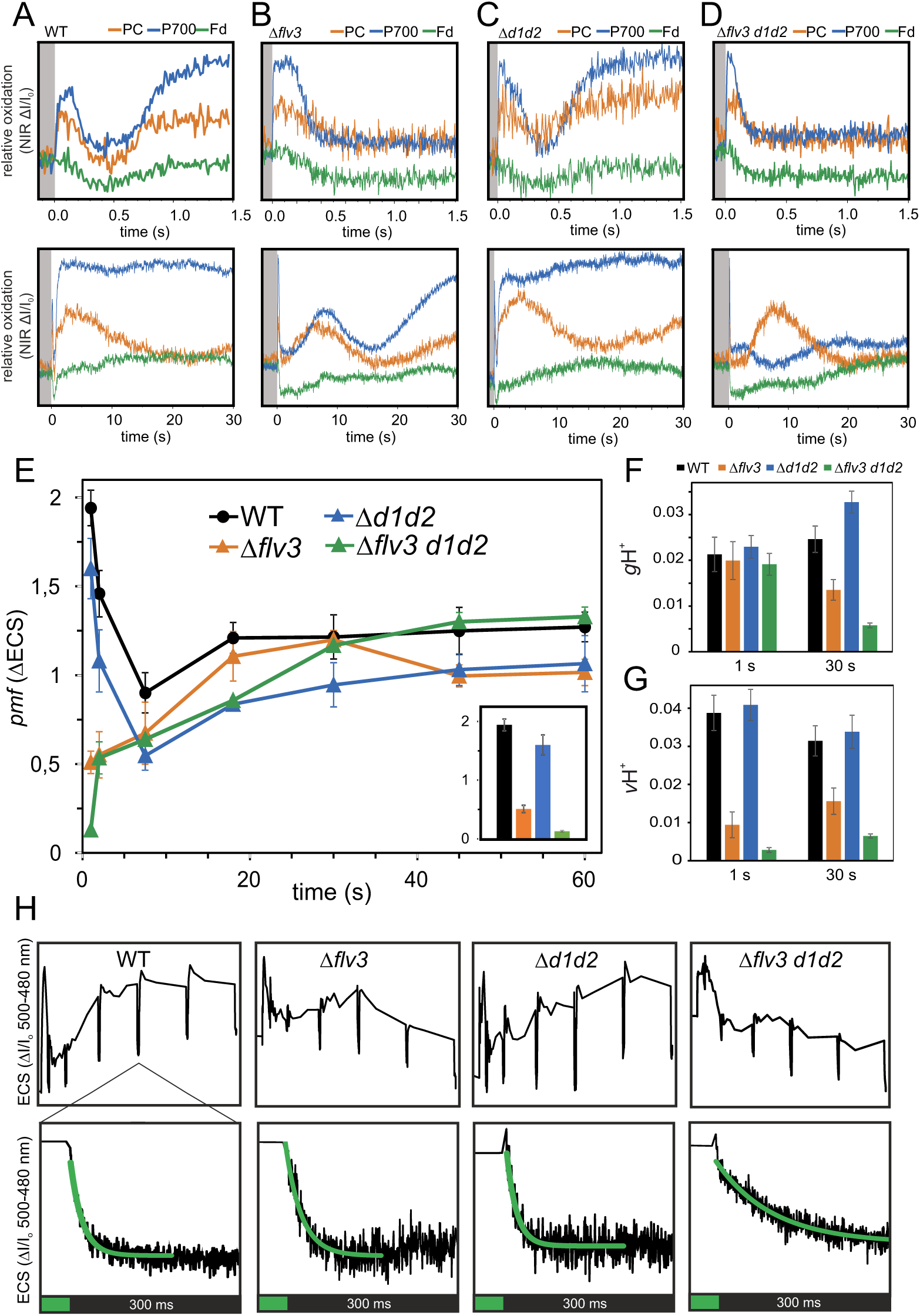
Redox changes of PC, P700, and Fd, and buildup of *pmf* during dark-to-HL transitions. WT **(A)**, Δ*flv3* **(B)**, Δ*d1d2* **(C)**, and Δ*flv3 d1d2* cells **(D)** were dark-adapted for 10 min, after which absorbance difference changes at 780–800 nm, 820–870 nm, 840–965 nm and 870–965 nm were measured with the Klas-NIR 100 spectrometer during 30 s illumination at 503 μmol photons m^-2^s^-1^ of red actinic light. The PC, P700, and Fd signals were deconvoluted from the absorbance difference changes using differential model plots (DMPs) determined for this study (Fig. S3). The upper panels in A–D show magnifications of the first 1.5 seconds of illumination from the lower panels. The experiment was repeated with three independent biological replicates, of which representative traces are shown. See also Supplemental Fig. S3. **(E)** Buildup of *pmf* during transitions from dark to 500 μmol photons m^-2^s^-1^ of green actinic light. Absorbance difference between 500-480 nm (ECS) was measured with a JTS-10 spectrometer, and light-induced *pmf* was determined from the dark interval relaxation kinetics (DIRK) of the ECS signal. Means ±SEM from 3–4 independent experiments are shown. The small inset shows *pmf* values after 1s of illumination for a clearer view. **(F)** Conductivity of the thylakoid membrane (gH+) after one and 30 s of illumination as in (E). gH+ was determined as the inverse of the time constant of first-order post-illumination decay kinetics of the ECS signal. **(G)** Thylakoid proton flux (vH+), calculated as *pmf* x gH+. **(H)** Representative traces of 500-480 nm absorbance differences used to determine the values in (E) and post-illumination ECS decay kinetics during a 500 ms dark interval after 30 s in light (lower panels). First-order fits are shown in green. Cells, after growth in air [CO_2_] / 50 μmol photons m^-2^s^-1^ for 4 days, were harvested and Chl *a* concentration adjusted to 10 μg ml^-1^ with fresh BG-11 for A–D and to 7.5 μg ml^-1^ for E–J.

To inspect whether the phenotypes observed above depend on differential buildup or regulation of *pmf*, we measured *in vivo* changes in the *pmf* by monitoring the absorbance change difference between 500-480 nm, which constitutes the electrochromic shift (ECS) in *Synechocystis* (Viola et al., 2019). *In vivo* measurement of light-induced ECS revealed that after 1 s of illumination of dark-adapted WT cells with 500 μmol photons m^-2^s^-1^, high *pmf* level was transiently generated, followed by decline during subsequent seconds (Fig. 5E). After ∼10 s of illumination *pmf* again increased towards a steadier value (Fig. 5E). Congruently with the strong reduction of P700 and Fd (Fig. 5B), the initial *pmf* peak after the first second of illumination was heavily dependent on the presence of Flv1/3. In both Δ*flv3* and Δ*flv3 d1d2 pmf* and thylakoid proton flux (vH+) were lower than in WT after 1 s (Fig. 5G). In Δ*flv3 pmf* remained drastically lower than in WT during the first seconds of illumination, but differed only slightly from WT thereafter due to diminished conductivity of the thylakoid membrane (gH+) (Fig. 5F), which is mainly determined by the activity of the ATP synthase (Cruz et al., 2005; Viola et al., 2019). In Δ*d1d2* thylakoid proton flux was similar to WT (Fig. 5G), likely due to enhanced FDP activity (Fig. 2C) compensating for impaired CET and respiration. However, *pmf* remained lower than in WT throughout the 1 min experiment (Fig. 5E) because of elevated conductivity (Fig. 5F). In contrast, despite drastically lower proton flux (Fig. 5G), Δ*flv3 d1d2* cells maintained *pmf* close to WT level after the first seconds in light (Fig. 5E) due to diminished conductivity (Fig. 5F). This indicates that the increased acceptor side limitation of PSI in Δ*flv3 d1d2* in comparison to Δ*flv3* (Fig. 3A) is not caused by lack of photosynthetic control as a consequence of lowered *pmf* generation in CET. Based on comparison of ECS kinetics in single, double, and triple mutant cells studied here, we conclude that both Flv1/3 and NDH-1_1,2_ contribute to proton flux during transitions from dark to high irradiance. However, deficiency in NDH-1_1,2_ in Δ*d1d2* is mostly compensated by elevated FDP-activity in terms of proton flux, while ATP synthase activity is increased, possibly as a response to the delay in activation of carbon fixation (Fig. 2C). In contrast, impaired thylakoid proton flux in Δ*flv3* during dark-to-light transitions cannot be compensated by NDH-1. Instead, downregulation of ATP-synthase activity lowers thylakoid conductivity and allows maintenance of *pmf* (apart from the first seconds of illumination) in Δ*flv3* and, even more dramatically, in Δ*flv3 d1d2*.

### PSI content is diminished in triple mutants deficient in Flv1/3 and NDH-1_1,2_ shifted to air [CO_2_] and high irradiance

The results from the growth assays, along with the real-time gas exchange, P700 redox change, and ECS measurements suggested that in Δ*d1d2* cells, an increase in the activity of FDPs compensates for the absence of a functional NDH-1_1,2_ complex, allowing efficient oxidation of PSI under high irradiance. Conversely, in Δ*flv1* and Δ*flv3* mutants, over-reduction of the electron transport chain occurs initially upon exposure to high irradiance, but another mechanism(s) eventually allow acclimation and survival in high [CO_2_] and high light (Helman et al., 2003). Under air [CO_2_] Flv1/3 hetero-oligomers catalyse transient O_2_ photoreduction, which is why no strong growth phenotype is observed in Δ*flv1* or Δ*flv3* under constant high irradiance (Allahverdiyeva et al. 2013, Santana-Sanchez et al. 2019). However, when both Flv1/3 and NDH-1_1,2_ are absent and the electron sink capacity of the cytosol is not elevated by high [CO_2_], cells are unable to oxidize PSI in high light, possibly resulting in lethal photodamage to PSI. To assess this hypothesis, we employed immunoblotting to provide estimates of the protein content of PSII (D1) and PSI (PsaB) subunits, orange carotenoid protein (OCP), large isoform of the ferredoxin-NADP+ oxidoreductase (FNR_L_), FDPs, as well as NdhD3 and the bicarbonate transporter SbtA after 24 hours exposure to different growth conditions.

WT and Δ*flv1* or Δ*flv3* cells grown at 3% [CO_2_] / ML and shifted to air [CO_2_] / ML at OD=0.1, accumulated similar levels of the PSII reaction center protein D1 and the PSI subunit PsaB. However, we detected a moderate decrease in the amount of both D1 and PsaB in the Δ*flv1 d1d2* and Δ*flv3 d1d2* triple mutants, while an increased amount of both proteins was detected in Δ*d1d2* (Fig. 6A). After 24 hours exposure to air [CO_2_] / HL, PsaB amount had decreased even further in Δ*flv1 d1d2* and Δ*flv3 d1d2* (Fig. 6C). Furthermore, 77K fluorescence spectra revealed that while Δ*flv3 d1d2* and WT cells grown under air [CO_2_] / ML had similar PSI/PSII ratios (Fig. 6D), the 24-hour exposure to air [CO_2_] / HL caused a dramatic decrease in the relative PSI fluorescence cross section (Fig. 6E). This strongly supports the hypothesis that loss of PSI contributes to the lethality ofthe shift to air [CO_2_] / HL. Interestingly, a substantial decrease in D1 content in air [CO_2_] / HL was also observed in the triple mutants (Fig. 6B-C).

**Figure 6.**
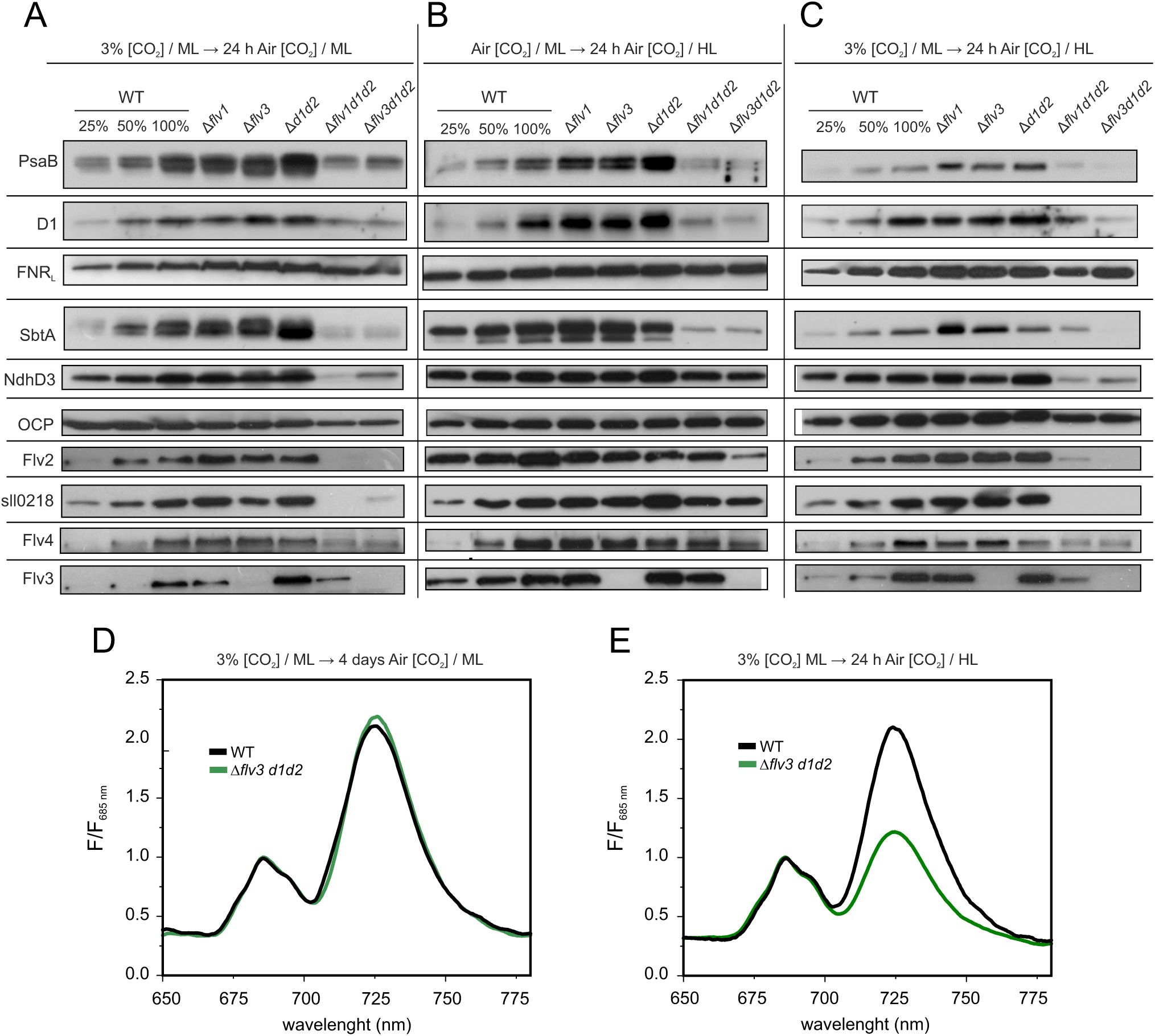
Immunodetection of proteins related to photosynthesis and CCM and 77K fluorescence emission spectra in WT and mutant strains. Pre-cultures were grown for 3 days at 3% [CO_2_] / ML (50 μmol photons m^-2^s^-1^) and then pelleted, resuspended in fresh BG-11 at OD_750_=0.15 and (**A**) incubated for 24 hours in air [CO_2_] / ML, (**B**) grown for 4 days in air [CO_2_] / ML, harvested, resuspended in BG-11 at OD_750_=0.15 and then incubated for 24 hours in air [CO_2_] / HL (220 μmol photons m^-2^s^-1^), (**C**) incubated for 24 hours in air [CO_2_] / HL. Total protein extracts were separated by SDS-PAGE and probed with specific antibodies. The immunoblots shown are representative of 2–3 independent biological replicates. (**D**) 77K fluorescence emission spectra from intact WT and Δ*flv3 d1d2* cells grown for 4 days in air [CO_2_] / ML. (**E**) After pre-growth at 3% [CO_2_] / ML cells were pelleted, resuspended in fresh BG-11 at OD_750_=0.15 and incubated for 24 hours in air [CO_2_] / HL. The cells (**D** and **E**) were pelleted, adjusted to [Chl *a*] of 7.5 μg ml^-1^, and excited with 440 nm light at 77K. The spectra were normalized to the PSII fluorescence peak at 685 nm.

Importantly, the Δ*flv1 d1d2* and Δ*flv3 d1d2* strains were unable to induce substantial accumulation of the proteins encoded by the *flv4-2* operon after a shift from 3% to air [CO_2_] (Fig. 6A and Fig. 6C). Only when pre-grown for 4 days in air [CO_2_] / ML before being shifted to higher irradiance, the expression of the operon was induced (Fig. 6B). Similarly to *flv4-2* operon proteins, both NdhD3 and SbtA failed to accumulate after shifts from 3% to air [CO_2_] / HL in the triple mutants, which likely impairs CCM and contributes to the lethal phenotype of the triple mutants in those conditions. Only small amounts of NdhD3 and SbtA were detected also in triple mutant cells shifted from 3% [CO_2_] to air [CO_2_] / ML (Fig. 6A). Closer to WT amount of NdhD3 but only a small amount of SbtA was detected after 24 hours in air [CO_2_] / HL (shifted from air [CO_2_] / ML) (Fig. 6B). In the Δ*d1d2* mutant NdhD3 content was similar to WT, but Flv3 as well as the *flv4-2* operon proteins (Flv2, Flv4, and Sll0218) were upregulated in all conditions tested (Fig. 6), which likely contributes to the increased rate of O_2_ photoreduction in that strain (Fig. 2C). The amounts of OCP and of FNR_L_ were unchanged in the mutant strains in all conditions (Fig. 6).

## DISCUSSION

Flv1/3 hetero-oligomers have been shown to be essential for photoreduction of O_2_ as a rapid response to excessive reduction of PSI, and consequently, for cell survival under fluctuating light conditions (Allahverdiyeva et al., 2013; Santana-Sanchez et al., 2019). Moreover, we recently showed that in air [CO_2_], Flv2/4 hetero-oligomers also catalyze low level light-induced steady-state reduction of O_2_ on the acceptor side of PSI in a coordinated manner with Flv1/3 (Santana-Sanchez et al., 2019).

In turn, another alternative electron transport component, the NDH-1 complex, functions in CET around PSI, using Fd as electron donor (Saura and Kaila, 2019; Schuller et al., 2019). Besides CET, the NDH-1_1,2_ complexes, incorporating the NdhD1 or NdhD2 subunit, are involved in respiration, whereas NDH-1_3,4_ complexes, incorporating the NdhD3 or NdhD4 subunit, function in CO_2_ uptake. In the current study, we have used *Synechocystis* mutant strains with different combinations of deficiencies in Flv1/3 and NDH-1 in order to examine the physiological significance of these alternative electron transport pathways and the possibility of functional interdependence between them. Our results provide compelling evidence for the concomitant physiological functions of Flv1/3 and NDH-1_1,2_ and their dynamic coordination for the efficient oxidation of PSI (thus protecting it from photodamage) under variable light conditions, and when low [CO_2_] limits the consumption of reductants in the CBB cycle. The two pathways can compensate for each other’s absence to some extent, but absence of both the Flv1/3 hetero-oligomer and NDH-1_1,2_ is lethal when cells are transferred from elevated [CO_2_] to the combined condition of air level [CO_2_] and high irradiance (Fig. 1D-E). In addition to the absence of these two important pathways, lethality is due to an inability to accumulate low C_i_-inducible photoprotective CCM proteins (Fig. 6C). Combined deficiency of Flv1/3 and NDH-1_3,4_ did not result in a lethal phenotype upon similar shifts, indicating that functional redundancy exists specifically between the Mehler-like reaction and NDH-1_1,2_

### Coordinated functions of Flv1/3 and NDH-1_1,2_ protect PSI by maintaining redox poise between the PETC and carbon fixation

Cells lacking functional Flv1/3 hetero-oligomers suffer from a transient over-reduction of PSI during sudden increases in light intensity (Fig. 3 and Fig. 5) due to the impairment of the Mehler-like reaction (Fig. 2, Allahverdiyeva et al., 2013). Whilst NAD(P)H has been proposed as the electron donor to Flv3 (Vicente et al., 2002; Brown et al., 2019) and Flv1 homo-oligomers (Brown et al., 2019), *in vivo* experiements have been unsupportive (Mustila et al. 2016), with exact electron donor(s) to FDP hetero-oligomers yet to be proven. *In vivo* experiments performed in this study demonstrate that the quick re-oxidation of Fd after 0.5 s (Fig. 5A) in light is absent in the Δ*flv3* deletion strains, whereby Fd remains strongly reduced (Fig. 5B). This result, together with a lack of impairment in NADP^+^ reduction and oxidation kinetics of Δ*flv3* deletion strains (similar conditions, Fig. 4F) provide strong support for Fd, rather than NAD(P)H, being the primary electron donor to the Flv1/3 hetero-oligomer *in vivo*. Accordingly, Fd has been shown to interact with Flv1 and Flv3 by Fd-chromatography (Hanke et al., 2011) and with Flv3 by a two-hybrid assay (Cassier-Chauvat and Chauvat, 2014).

The deficiency of NDH-1_1,2_, in turn, reduced the cells’ ability to maintain oxidized P700 and Fd under high irradiance soon after (∼50–200 ms) a dark-to-light transition (Fig. 5C). This brief timescale may be due to PSI-NDH-1 supercomplexes (Gao et al., 2016) where P700 can be oxidized rapidly upon the onset of illumination. It has been shown that formation of the NDH-1-PSI supercomplex is important to keep PSI functional under various stress conditions (Zhao et al. 2017). NDH-1_1,2_ -deficiency also caused slightly elevated acceptor side limitation of PSI under low irradiance (Fig. 3A), which was likely due to a lack of NADPH oxidation during the low light phases of fluctuating light (Fig. 4C). Thus, NDH-1_1,2_ appears to play an important role under low light, as has been previously suggested for chloroplastic NDH in angiosperms (Yamori et al., 2011; Yamori et al., 2015) and bryophytes (Ueda et al., 2012). The delayed activation of CBB in Δ*d1d2* did not result in an inability to oxidize PSI (Fig. 3B and Fig. 5C), likely due to significant enhancement of FDP-mediated O_2_ photoreduction (Fig. 2C) providing an enlarged electron sink for the photosynthetic electron transport chain.An opposite order of causation is also possible, whereby the excessive funneling of photosynthetic electrons to O_2_ would cause the delay in induction of CO_2_ fixation in Δ*d1d2*. However, the observation that there is no NADPH shortage at the onset of illumination in Δ*d1d2,* and rather the consumption of NADPH during transitions from HL to LL is impaired (Fig. 4C), suggests that the availability of reductant for the CBB is not the limiting factor. Nor is it likely CO_2_, as CCM is functioning as in WT (Fig. 2C), or ATP, as ATP synthase activity was even higher than in WT in Δ*d1d2* (Fig. 5F). Nevertheless, simultaneous deficiency of FDPs in addition to NDH-1_1,2_ (in Δ*flv3 d1d2* triple mutant), with the effect of diminishing the flow of photosynthetic electrons to O_2_ photoreduction, mostly rescued the delay in CO_2_ fixation (Fig. 2D) as well as NADPH consumption (Fig. 4D) seen in Δ*d1d2*. Transient O_2_ photoreduction at a low rate was still observed in Δ*flv3 d1d2* during dark-to-HL transitions (Fig. 2D), possibly mediated by the thylakoid terminal oxidases (Ermakova et al., 2016) or by photorespiration (Allahverdiyeva et al., 2011). However, the triple mutants suffered from more severe inability to oxidize PSI than Δ*flv3* during sudden increases in irradiance (Fig. 3). These observations suggest that in addition Flv1/3, also NDH-1_1,2_ has a role in contributing to oxidation of PSI during changes in light conditions or carbon availability. Upon deficiency of NDH-1_1,2_ (in Δ*d1d2*), cells prioritize protection of PSI over efficient CO_2_ fixation by upregulating the Mehler-like reaction via an unknown mechanism. The triple mutants cannot do this, leading to timely induction of CO_2_ fixation at the high cost of inability to oxidize PSI (Fig. 3A and 3) or Fd (Fig. 5D). This results in loss of PSI (Fig. 6), possibly due to photodamage to its FeS clusters (Tiwari et al., 2016; Shimakawa et al., 2016).

### On the mechanism of NDH-1-mediated oxidation of PSI

Contribution of NDH-1 to oxidation of PSI during sudden increases in light intensity is not unprecedented. In angiosperms, where FDPs have been lost during evolution (Ilik et al., 2017), *Arabidopsis* mutants lacking the NDH complex show a delay in oxidation of PSI during increases in light intensity in comparison to *Arabidopsis* WT (Nikkanen et al., 2018; Shimakawa and Miyake, 2018a). By definition, canonical CET cannot directly increase the relative proportion of oxidized P700, as electrons from the acceptor side of PSI are shunted back to the intersystem electron transfer chain. However, NDH-1 may enhance PSI oxidation by at least three alternative, but mutually non-exclusive mechanisms.

i) NDH-1-mediated CET is coupled to translocation of protons from cytosol to the thylakoid lumen with a 2H^+^/e^-^ stoichiometry (Strand et al., 2017; Saura and Kaila, 2019). It will therefore contribute to buildup of ΔpH, which limits electron transfer to PSI by inhibiting PQH_2_ oxidation at Cyt *b*6*f* (Shimakawa and Miyake, 2018b) and drives ATP synthesis to accommodate the needs of the CBB, increasing its electron sink capacity. As the Mehler-like reaction also contributes to buildup of ΔpH by consuming H^+^ on the cytosolic side of the thylakoid membrane and by supporting linear electron flow (Fig. 5), (Allahverdiyeva et al., 2013), enhanced FDP activity in Δ*d1d2* (Fig. 2C) partly compensates for the lack of NDH-1_1,2_ in respect to generation of proton flux (Fig. 5G). However, this fails to explain the exacerbated impairment of P700 and Fd oxidation in Δ*flv3 d1d2* in comparison to Δ*flv3* (Fig. 3 and Fig. 5) because in the triple mutant *pmf* was not lower than in Δ*flv3* (Fig. 5E). Adequate *pmf* is maintained however at the expense of ATP synthase activity (Fig. 5F), and it is likely that diminished ATP production contributes to the increased acceptor side limitation of PSI in Δ*flv3 d1d2* (Fig. 3A). NDH-1_1,2_ could therefore contribute to P700 oxidation by enhancing cytosolic sink capacity by providing a more suitable ATP:NADPH ratio for the CBB. Adjustment of the ATP:NADPH ratio closer to the theoretically optimal 3:2 has long been considered a fundamental reason for the existence of CET (Kramer et al., 2004; Yamori and Shikanai, 2016).
ii) NDH-1-mediated respiratory electron transfer, *i*.*e*. coupling of PQ reduction by NDH-1 to O_2_ reduction by thylakoid terminal oxidases (*i*.*e*. Cyd and Cox) would also contribute to oxidation of PSI by relieving electron pressure in the intersystem chain during illumination (Ermakova et al., 2016), as well as further contribute to ΔpH by consuming H^+^ on the cytosolic side of thylakoids (Cyd) and pumping protons to the lumen (Cox) (Brändén et al., 2006). Interestingly, Liu and colleagues (2012) have shown that the subcellular localization of NDH-1 complexes is dependent on the redox state of the PQ pool. An oxidized PQ pool causes NDH-1 to accumulate at specific clusters in thylakoid membranes where it would be more likely to transfer electrons (via the PQ pool) to a terminal oxidase. A reduced PQ pool, in turn, results in a more even distribution of NDH-1 within thylakoids (Liu et al., 2012). However, it is important to note that terminal oxidases do not have a high electron sink capacity (Ermakova et al., 2016).
iii) NDH-1 has been predicted, albeit not yet experimentally shown in photosynthetic organisms, to be able to function in reverse: to oxidize PQH_2_ driven by concomitant release of protons from the thylakoid lumen (Strand et al., 2017). Such reverse activity would constitute a ‘pseudo-linear’ electron transfer pathway that would bypass PSI and thereby prevent its over-reduction. This could occur in conditions where the PQ pool is reduced, *pmf* is high, and the Fd pool is oxidized (Strand et al., 2017). Such conditions likely exist transiently during dark-to-light and LL-to-HL transitions (Fig. 4 and 5) (Strand et al., 2019). Accordingly, fast re-reduction of P700^+^ already after 50 ms was observed in Δ*d1d2* and Δ*flv3 d1d2* during dark-to-HL transitions (Fig. 5C). As supported by the impaired ability to oxidize Fd in absence of Flv1/3 (Fig. 5), the presence of Flv1/3-catalyzed Mehler-like reaction would likely be essential in this model in order to maintain the Fd pool in a sufficiently oxidized state to provide electron acceptors for reverse-functioning NDH-1. It is noteworthy, however, that NDH-1 reverse activity would also have the effect of lowering ΔpH, thereby relieving photosynthetic control at Cyt *b*6*f*. This could counteract the effect of any reverse NDH-1 activity by increasing electron flow to PSI.

Hypothetical mechanisms for coordination of Flv1/3 and NDH-1_1,2_ activities are shown in Fig. 7.

**Figure 7.**
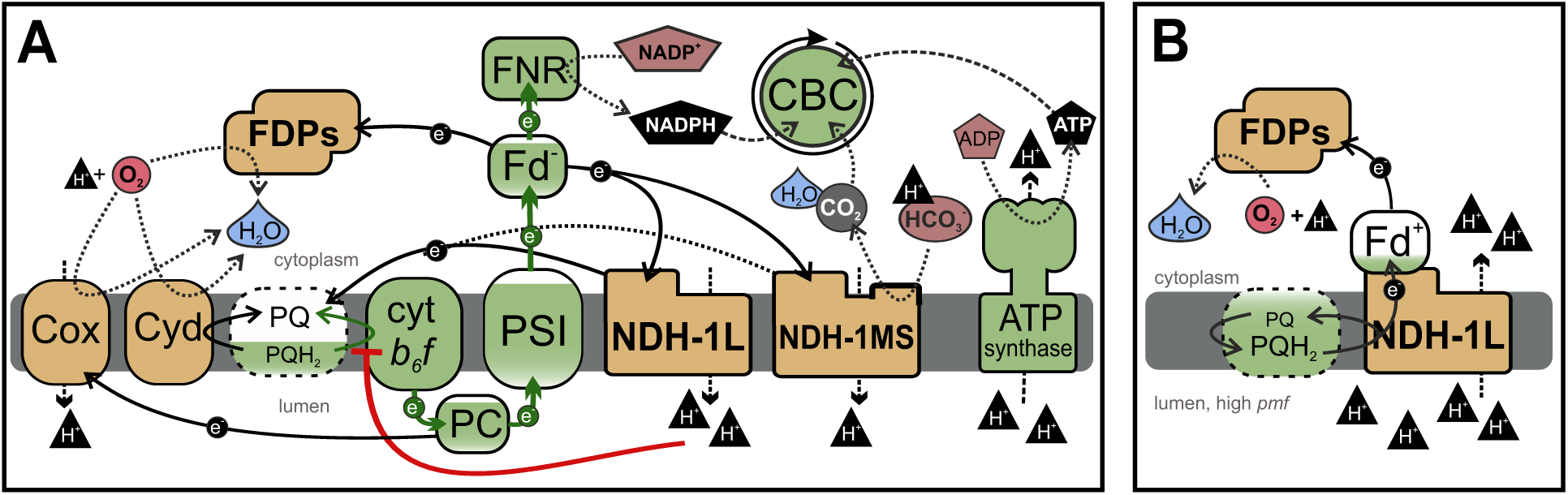
Schematic models for the coordination of the activities of FDPs and NDH-1 in preventing over-reduction of PSI. (**A**) In conditions where the Fd pool is higly reduced and the PQ pool is not, NDH-1L transfers electrons from Fd to the PQ pool, coupled to O_2_ reduction by thylakoid-localised RTOs. NDH-1L pumps 2H^+^/e^-^ into the lumen while FDPs and RTOs consume H^+^ when reducing O_2_ to H_2_O on the cytosolic side of thylakoids. This increases transthylakoid *pmf*, which in turn drives ATP synthesis as well as inhibits PQH_2_ oxidation at Cyt *b*_6_*f*, thereby preventing excessive electron supply to PSI. Carbonic anhydrase activity of NDH1-MS converts HCO_3_^-^ to CO_2_, consuming H^+^ in the cytosol, likely also coupled to translocation of H^+^ to the lumen, further increasing *pmf*. Increased availability of CO_2_ and ATP/NADPH ratio enhances the electron sink capacity of the CBB on the acceptor side of the PETC. (**B**) If the PQ pool is highly reduced, Fd pool is oxidized, and *pmf* is high, NDH-1L could hypothetically function in reverse, transferring electrons from PQH_2_ to Fd^+^, powered by release of 2H^+^/e^-^ from the lumen. A strong Mehler-like reaction catalyzed by Flv1/3 would be required to maintain a sufficient pool of oxidized Fd, with Flv1/3 likely accepting electron directly from Fd^-^. Most components depicted in (A) are omitted from (B) to enhance clarity, not to suggest their absence. Extent of the colour fill for PQ/PQH_2_ and Fd depict level of reduction.

### Inability to induce a strong CCM network contributes to lethality upon shifts to air [CO_2_] / high irradiance in cells deficient in both Flv1/3 and NDH-1_1,2_

Exposure of *Synechocystis* cells to low [CO_2_] induces expression of high-affinity CCM-related genes such as SbtA, a HCO_3_^-^ / Na^+^ symporter on the plasma membrane, and NdhD3, which is part of the NDH-1_3_ complex on thylakoids specializing in C_i_ uptake (Ohkawa et al., 2000; Shibata et al., 2001, Shibata et al., 2002; Zhang et al., 2004). CCM is energetically expensive, but the large C_i_ flux involved in the operation of CCM contributes to dissipation of excess light energy under stress conditions (Xu et al., 2008; Burnap et al. 2015). NDH-1_3,4_ in particular has a photoprotective role, using reduced Fd to drive CO_2_ conversion to HCO_3_^-^ with concomitant translocation of protons into the thylakoid lumen.

Despite the low induction level of the CCM proteins NdhD3 and SbtA, the triple mutants (Fig. 6) survive under standard growth conditions of air [CO_2_] / ML (Fig. 1B-C). When a decrease in [CO_2_] is coupled with an increase in irradiance, however, a low amount of CCM, reflected by low accumulation level of NdhD3 and SbtA in the Δ*flv3d1d2* and Δ*flv3d1d2* triple mutants (Fig. 6C) fails to dissipate excess energy, thus resulting in photodamage and lethality of those conditions. The regulatory mechanisms controlling the inability of the triple mutants to accumulate low Ci-inducible proteins is unclear and remains to be elucidated. Accumulation of NADP^+^ and α-ketogutarate inhibits the induction of CCM gene expression via interaction with the transcription factor NdhR (CcmR) (Daley et al., 2012). NdhR also controls expression of the Flv4-2 operon encoding Flv2, Flv4, and Sll0218 (Eisenhut et al., 2012), all of which also showed decreased accumulation levels in the triple mutants (Fig. 6). However, at least in the tested conditions, no substantial increase in the relative amount of NADP^+^ was detected in Δ*flv3d1d2* cells (Fig. 4). It is important to note however, that Flv1/3 and NDH-1_3,4_ cooperation alone is not crucial for cell metabolism, since the Δ*flv1 d3d4* and Δ*flv3 d3d4* mutant cells do not show clear growth phenotypes (Fig. S1).

## CONCLUSIONS

In the current study we have shown that FDPs and NDH-1 function cooperatively to maintain redox balance between the PETC and cytosolic carbon assimilation upon sudden changes in light intensity and/or carbon availability. It is likely that both pathways receive electrons primarily from Fd, which enables coordinated regulation of their activities. Cooperation of FDPs and NDH-1 has an essential photoprotective role during shifts to air [CO_2_] /HL, contributing to efficient oxidation of PSI, buildup of *pmf,* and induction of expression of low C_i_-specific CCM-related genes. Finally, it is worth noting that simultaneous removal of Flv1/3 and NDH-1_1,2_ as competitive electron sinks downstream of PSI may be a useful biotechnological tool for maximizing direction of photosynthetic electrons to desired pathways under controlled environmental conditions (McCormick et al., 2013; Thiel et al., 2019; Jokel et al., 2019). Furthermore, our findings about functional interplay between FDPs and NDH-1 will be highly relevant in projects aiming to enhance crop productivity via introduction of exogenous FDPs to higher plants (Yamamoto et al., 2016; Gómez et al., 2018).

## MATERIALS AND METHODS

### Strains and culture conditions

In the current study we used the glucose-tolerant WT strain *Synechocystis* sp. PCC 6803 (Williams, 1988), single mutant strains Δ*flv1* and Δ*flv3* (Helman et al., 2003), the M55 mutant (Δ*ndhB*) (Ogawa 1991), double mutants Δ*d1d2 and* Δ*d3d4* (Ohkawa et al., 2000a), and triple mutants Δ*flv1 d1d2,* Δ*flv3 d1d2*, Δ*flv1 d3d4,* and Δ*flv3 d3d4* obtained from the CyanoMutants collection (Nakamura et al., 1999). The triple mutants were constructed by T. Ogawa in Δ*d1d2 and* Δ*d3d4* backgrounds. All the mutants demonstrated complete segregation. Pre-experimental cultures were always grown in 30 ml batches of BG-11 medium pH 7.5 (Williams, 1988) under 3% [CO_2_] at 30°C under continuous white light of 50 µmol photons m^−2^s^−1^ (ML) with agitation. Mutant pre-cultures were supplemented with the appropriate antibiotics. At the logarithmic growth phase, cells were harvested and re-suspended in fresh BG-11 without antibiotics at OD_750_ of 0.1–0.2, as described in the appropriate figure legends. Cells were then shifted to air [CO_2_] / 30°C and illuminated continuously with white light of either 50 or 220 µmol photons m^−2^s^−1^, as described in the figure legends.

### Gas exchange measurements

Exchange of ^16^O_2_ (*m/z*=32), ^18^O_2_ (*m/z*=36), CO_2_ (*m/z*=44) was measured *in vivo* with membrane-inlet mass spectrometry (MIMS) as described in Mustila et al. (2016). Harvested cells were resuspended in fresh BG-11 pH 7.5 and adjusted to 10 µg Chlorophyll (Chl) *a* ml^−1^, and kept for one hour in air [CO_2_] / 50 µmol photons m^−2^s^−1^. Prior to the measurements, cells were supplemented with ^18^O_2_ at an equivalent concentration to ^16^O_2_ and with 1.5 mM NaHCO_3_. Cells were dark-adapted for 15 min, after which gas exchange was monitored over a 5 min illumination period of 500 μmol photons m^-2^s^-1^ of white actinic light (AL). The gas exchange rates were calculated according to Beckmann et al., (2009).

### Measurement of fluorescence and absorbance changes

Experimental cultures for all spectroscopic experiments were grown for four days in air [CO_2_] / 50 µmol photons m^−2^s^−1^ at 30°C in BG-11 pH 7.5. Chl *a* fluorescence and P700-oxido-reduction (875-830 nm absorbance difference) were simultaneously recorded (Fig. 3) with the Dual-PAM 100 spectrophotometer (Walz, Effeltrich, Germany). Harvested cells were resuspended in fresh BG-11 pH 7.5 and adjusted to 10 µg Chl *a* ml^−1^, and kept for one hour in air [CO_2_] / 50 µmol photons m^−2^s^−1^ and dark-adapted for 10 min before being subjected to fluctuating light regime alternating between 1 min periods of 25 and 530 µmol photons m^−2^s^−1^ of red AL. Saturating pulses (500 ms of 5000 μmol photons m^-2^s^-1^) were administered at 15 s intervals. Photosynthetic parameters were calculated as follows: Y(NA)=(P_m_-P_m_’)/P_m_; Y(ND)=P/P_m;_ Y(I)=1-Y(ND)-Y(NA) (Klughammer and Schreiber, 2008); Y(II)=(F_m_’-F)/F_m_’ (Genty et al., 1989).

A Dual-Klas NIR 100 spectrophotometer (Walz) was used to measure absorbance difference changes at 780–800 nm, 820–870 nm, 840–965 nm and 870–965 nm, from which PC, P700, and Fd redox changes (Fig. 5) were deconvoluted based on differential model plots (DMPs) for PC, P700, and Fd (Klughammer and Schreiber, 2016; Schreiber, 2017; Setif et al., 2019). In WT *Synechocystis*, very fast re-oxidation of Fd upon illumination impedes measurement of model spectra. To circumvent this issue, we measured the DMPs from Δ*flv3 d1d2* cells where P700 oxidation was severely delayed in comparison to WT (Fig. 3B). The DMPs (Fig. S3) were measured using scripts provided with the instrument software. Harvested cells were adjusted to 10 µg Chl *a* ml^−1^ with fresh BG-11 pH 7.5, and kept for one hour in air [CO_2_] / 50 µmol photons m^−2^s^−1^ and dark-adapted for 30 min before illumination for 30 s with 503 μmol photons m^-2^s^-1^ of red AL. It is noteworthy that cytochrome *c_6_* and (F_A_F_B_) redox changes likely contribute to the PC and Fd signals, respectively (Setif et al., 2019).

NADPH fluorescence changes between 420 and 580 nm, induced by excitation at 365 nm, were measured simultaneously with Chl *a* fluorescence with the Dual-PAM 100 and its 9-AA/NADPH accessory module (Walz) (Schreiber and Klughammer, 2009; Kauny and Setif, 2014). Experimental samples were prepared as above and [Chl *a*] was adjusted to 5 µg ml^−1^. Cells were dark-adapted for 10 min and subjected to the same fluctuating light regime as in Fig. 3, but without saturating pulses.

Electrochromic shift (ECS) was measured as absorbance difference between 500 and 480 nm (Viola et al., 2019). Absorbance changes were measured from dark-adapted cell suspensions with a JTS-10 spectrophotometer (BioLogic, Seyssinet-Pariset, France) using 500 and 480 nm CWL 50mm 10nm FWHM bandpass filters (Edmund Optics, Barrington, NJ, USA) and BG39 filters (Schott, Mainz, Germany) protecting the light detectors from scattering effects. Absorbance changes induced by measuring light only were subtracted from changes under ML + AL. ECS values were normalized to ECS change induced by a single turnover flash provided by a XST-103 xenon lamp (Walz). Experimental samples were prepared as above and [Chl *a*] was adjusted to 7.5 µg ml^−1^, and illuminated with green AL of 500 μmol photons m^-2^s^-1^ for 1s or 60 s interspersed with 500 ms dark intervals at 2, 7.5, 18, 30, and 45 s. *Pmf* was calculated as the extent of ECS decrease at the dark intervals. Thylakoid conductivity (gH+) was calculated as the inverse of the time constant of a first-order fit to ECS relaxation kinetics during a dark interval, and proton flux (vH+) as *pmf* x gH+ (Cruz. et al., 2005).

77K fluorescence emission spectra were measured with a QE Pro spectrophotometer (Ocean Optics, Dunedin, FL, USA). Cells were harvested and [Chl *a*] was adjusted to 7.5 µg ml^−1^ with fresh BG-11 pH 7.5. Cells were frozen in liquid N_2_ and excited at 440 nm. Raw spectra were normalized to PSII-fluorescence peak at 685 nm.

### Protein extraction and immunoblotting

Total protein extracts from cultures were isolated as previously described (Zhang et al., 2009), separated by SDS-PAGE on 12 % gels with 6M urea, and blotted on PVDF membranes.

Membranes were probed with antibodies raised against D1, PsaB (Agrisera, Vännäs, Sweden, AS10 695), PetH (Kindly shared by H. Matthijs), OCP (kindly shared by D. Kirilovsky), SbtA (kindly shared by T. Ogawa), NdhD3 (Eurogentec, Liége, Belgium), Flv2, Flv4, Flv3 (Antiprot, Puchheim, Germany), sll0218 (MedProbe, Oslo, Norway). Horseradish peroxidase –conjugated secondary antibody (GE Healthcare, Chicago, IL, USA) and Amersham ECL (GE Healthcare) were used for detection.

## Supplemental material

Supplemental Figure S1. Growth of Δ*flv1 d3d4* and Δ*flv3 d3d4* mutant strains.

Supplemental Figure S2. O_2_ and CO_2_ fluxes in M55 and in WT at pH 6.

Supplemental Figure S3. Near-infrared (NIR) differential model blots (DMPs) for deconvolution of PC, P700, and Fd signals with the Klas-NIR 100 spectrometer.

## ACKNOWLEDGEMENTS

We thank Prof. Teruo Ogawa for sharing the FDP and NDH-1 mutants and Dr. Henna Mustila for construction of the ΔM55 *flv3* strain. Dr. Duncan Fitzpatrick is thanked for expertise and assistance with MIMS. We also thank Dr. Pierre Setif and Dr. Gert Schansker for their expert advice on measuring the DMPs for the Dual-Klas NIR spectrophotometer, and Prof. Jens Appel for critical reading of the manuscript. We declare no conflicts of interest.

